# Ebola virus disease outbreak in the Republic of Guinea 2021: hypotheses of origin

**DOI:** 10.1101/2021.04.23.440924

**Authors:** A.A. Kritsky, S. Keita, N. Magassouba, Ya.M. Krasnov, V.A. Safronov, E.V. Naidenova, A.P. Shevtsova, E.A. Naryshkina, S.A. Shcherbakova, A.Yu. Popova, V.V. Kutyrev

## Abstract

In 2013-2016, a large-scale Ebola virus diseas (EVD) outbreak hit the countries of West Africa for the first time. Thus provoking a health crisis. The most affected countries were Sierra Leone, Liberia and the Republic of Guinea. Since the end of the outbreak in 2016, there have been no new cases until recently. In February 2021, the Republic of Guinea announced about laboratory-confirmed EVD cases in its territory.

By April 12, 2021, an Ebola virus disease outbreak in the province of N’Zerekore (Gueke sub-prefecture) of the Republic of Guinea had affected 16 people, 9 of whom had died. The origin of this outbreak is still unknown. This paper considers various hypotheses of its emergence. Within the frames of the study sequencing and analysis of the whole genome sequence of the strain that initiated the epidemic process have been carried out. The most likely cause of this outbreak, according to the results obtained, should be deemed a long-term persistence of the virus in the body of survivors, followed by a relapse of the disease and transmission among contact persons.

## Introduction

Ebola virus disease (EVD) – is a dangerous infectious disease, the etiological agent of which is the *Ebolavirus*. It is a viral hemorrhagic fever with a high letality rate (up to 90%). Outbreaks occur periodically in sub-Saharan African countries. The largest outbreak occurred in 2013-206 in West African countries, particularly affecting Sierra Leone, Liberia, and the Republic of Guinea. In total, over 28,000 cases were reported during that outbreak, more than 11,000 deaths. Since the end of the outbreak in 2016, West Africa has remained EVD-free. However, in February 2021, the Republic of Guinea reported new laboratory-confirmed EVD cases.

By April 12, 2021, an EVD outbreak in the province of N’Zerekore (Gueke sub-prefecture) of the Republic of Guinea had affected 16 people, 9 of whom had died. Laboratory confirmed cases were recorded between February 12 and April 2. 5 cases of EVD were registered among healthcare workers. The patients showed symptoms of diarrhea, vomiting and bleeding, which developed after attending a relative’s funeral on February 1, 2021. The origin of this outbreak remains unknown.

The aim of our study was to investigate the hypotheses about the most probable origin of the source of infection.

### Case description and hypothesis

The first known case is a female healthcare professional who initially contacted Gueke Medical Center on January18, 2021 complaining about headache, physical weakness, nausea, vomiting, loss of appetite, abdominal pain and fever, where she was mistakenly diagnosed with “typhoid” and then “malaria”. On January 24, against the background of a worsening condition, the patient turned to a traditional healer in N’Zerekore. On January 28, she died and was buried on February 1 without observing biological safety measures, which led to infection of the traditional healer and the relatives of hers [1]. The first laboratory confirmed cases of the disease among contact persons date back to February 12 and 13, 2021. After that, laboratory confirmed cases were reported almost daily from February 19 to February 27. The last laboratory confirmed case was registered on April 2. All of those cases were confined to N’zerekore Prefecture.

It should be noted that the clinical misdiagnosis at the first stage of the outbreak, the involvement of traditional healers and medical workers, as well as the pivotal role of unsafe burial rites, is the typical combination of signs of EVD epidemic process characteristic of West Africa, where the unprecedented epidemic was recorded in 2013-2016, which struck Guinea, Liberia and Sierra Leone (over 28 thousand cases, 11 thousand deaths), as well as, to a small extent, the countries of Europe and the United States [2].

To date, the cause of this epidemic outbreak remains unknown. As working hypotheses, the following can be formulated:

1. Import from the Central African region, where EVD cases have been reported in the DR Congo for quite a long time.
2. Latent chain of transmission of EVD among the population of the N’Zerekore region (Republic of Guinea).
3. Circulation of the EVD pathogen in a environmental with subsequent passage into the human population.
4. Transmission of the virus from survivors.

We examined a sample of the EVD agent genome isolated from clinical specimen (laboratory ID E2) obtained from a patient who had contact with the first case of EVD living in the N’Zerekore prefecture.

### Sample processing and sequencing

Extraction of RNA from a sample of biological material was performed using a kit for the extraction of nucleic acids “RiboPrep”, RNA detection was done using a RT - PCR kit «AmpliSens EBOV Zaire-FL», synthesis of cDNA on the RNA template was accomplished using the “Reverta-L” kit («InterLabService», Russia). Whole genome sequencing was performed on the IonS5TM XL System («ThermoFisher Scientific», USA). After sequencing, genome fragments were assembled into a single sequence aligning to the reference isolate Makona-Kissidougou-C15 (KJ660346, March 17, 2014, Guinea) in the Unipro UGENE v37 (http://ugene.net/) and MEGA 7.0 (http://www.megasoftware.net) software packages. To carry out a comparative analysis of our data, we used a representative sample from the international database NCBI GenBank (https://www.ncbi.nlm.nih.gov/genbank), containing 1814 genomes obtained from clinical specimens from patients with EVD in the countries of West and Central Africa (Republic of Guinea, Sierra Leone, Liberia, Mali, Nigeria, DR Congo, Gabon). MAFFT v.7 (https://mafft.cbrc.jp/alignment/server/) was used to align the genome sequence. The search for single nucleotide polymorphisms (SNPs) in the compared genomes was conducted using the Snippy 2.0 program (https://github.com/tseemann/snippy). Comparative phylogenetic analysis was performed using the BioNumerics 7.6 software package («Applied Maths NV», Belgium) and the Maximum Parsimony algorithm. The genome has been deposited in the international NCBI GenBank database under the access number: MW962392

### Genome analysis and hypothesis checking

Analysis of the obtained complete genome shows that the similarity of the genome fragments with ZE from West Africa (Liberia, Republic of Guinea, Sierra Leone) is 99,93%, while that with the genome of ZE from Central Africa (DRC, Gabon) is only 96-98%, which excludes the option of importing from the DR Congo. Strains from Central Africa form separate clusters distant from the cluster of isolates from West Africa on the phylogenetic tree (**Figure 1a**).

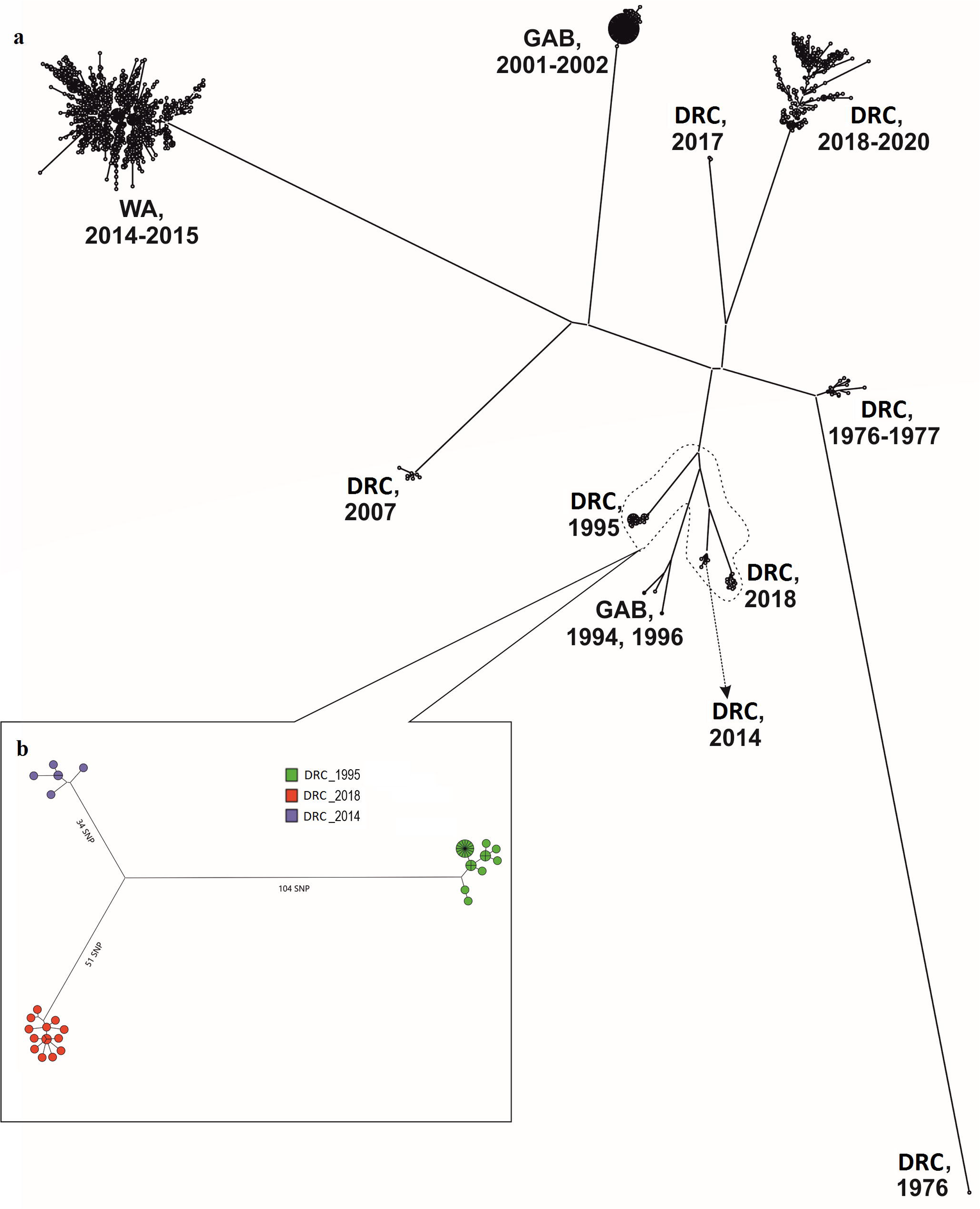

It follows from the data presented that the genetic profile of the EVD that caused the outbreak of 2021 practically does not differ from the strains found during the 2013–2016 outbreak in West Africa, forming a separate group [3] and is in the same phylogenetic cluster with them (**Figure 2**). At the same time, the strains from the Republic of Guinea and Liberia that circulated there in 2014 are the closest to the strain of 2021 (**Figure 3**). Thus, the difference in SNPs with the strains of 2014 is only 10 SNPs for the closest group of ZE strains from the Republic of Guinea and Liberia (MH425138, KT725387, KT725385, KT725263, KR817134, KR817132, KR817123, KR81745121) (**Figure 3**). The mutations found in the ZE genome from sample E2 relative to the genome of the reference strain Makona-Kissidougou-C15 (KJ660346, 17.03.2014, Guinea) are presented in the **table**.

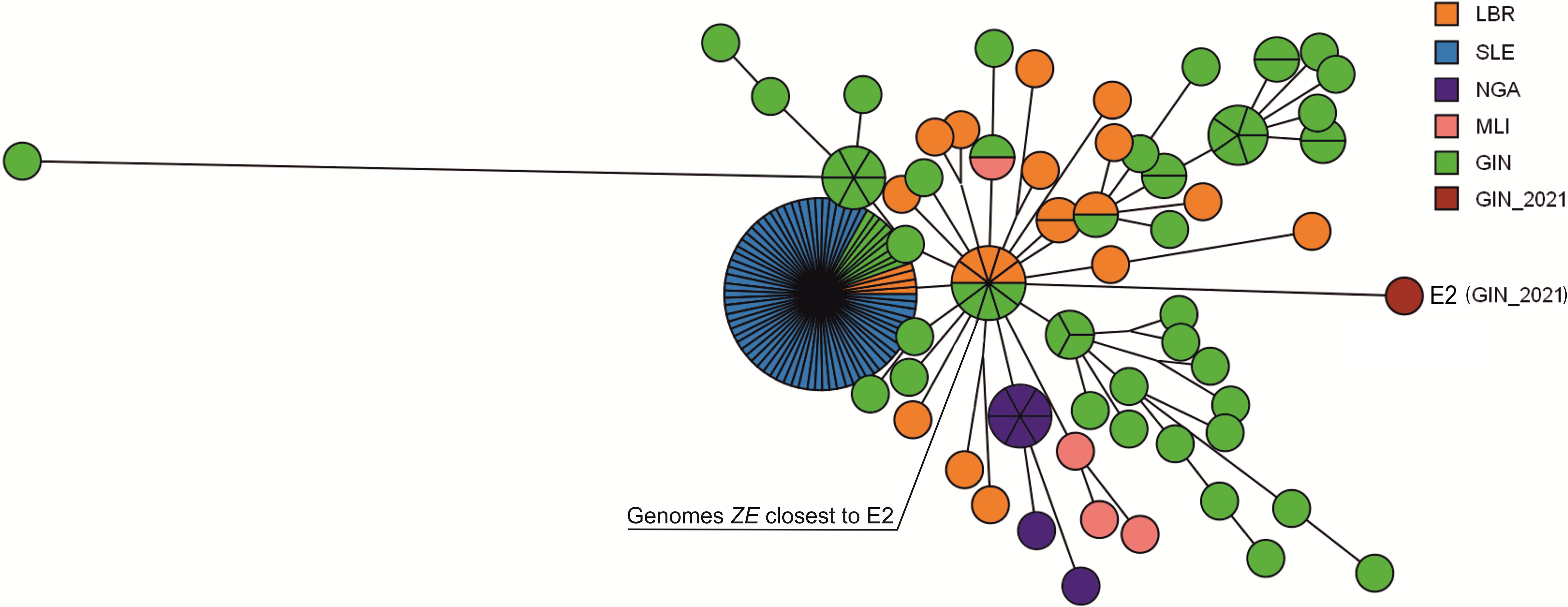

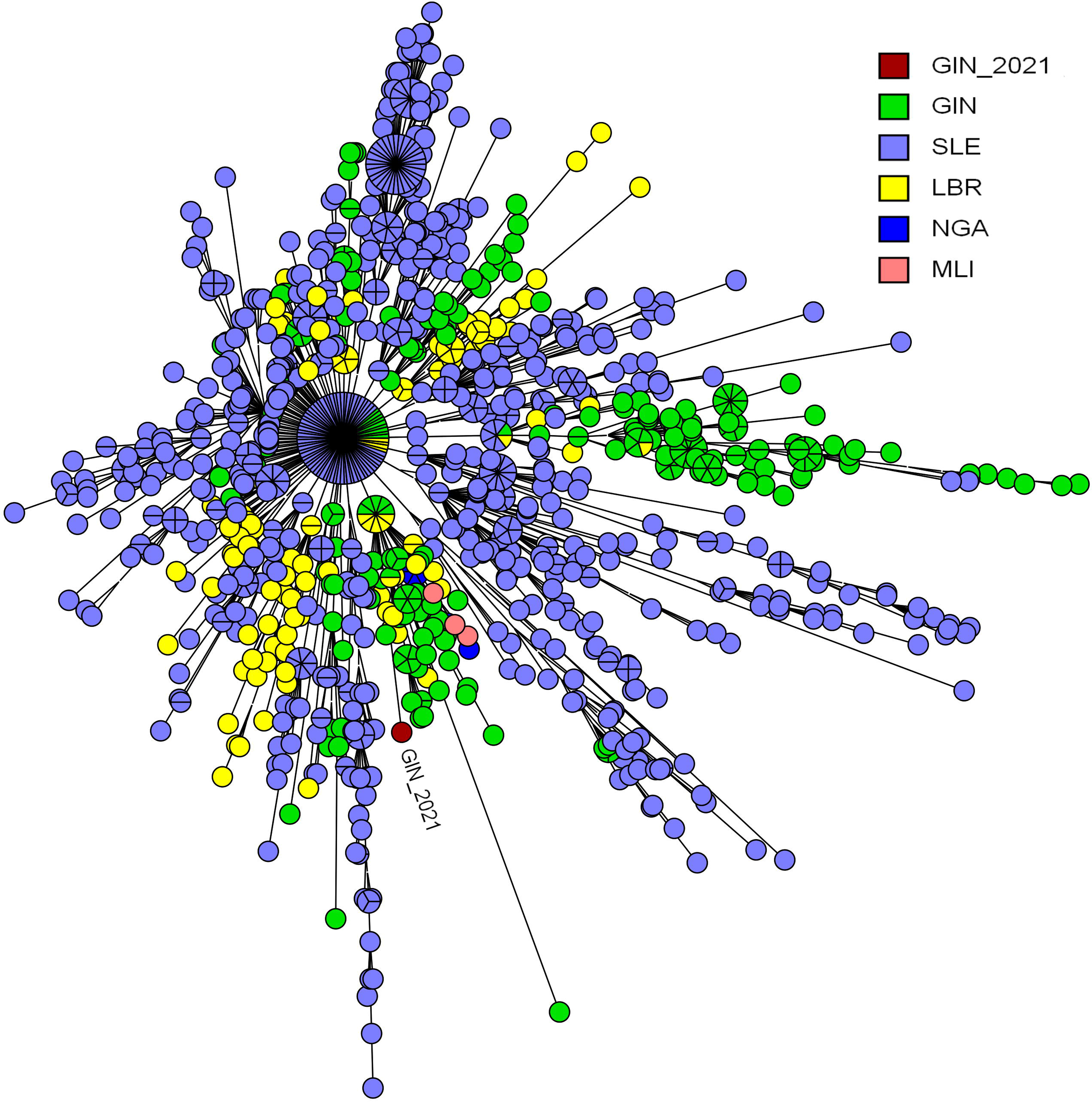

Taking into account the established frequency of SNPs occurence characteristic of ZE in case of person-to-person transmission (1–2×10^−3^ substitutions per site per year), it is not possible that new cases are the result of a latent chain of transmission among humans (there are 10 single substitutions instead of 100 or more expected in 5-6 years term) [4].

Frequency of occurrence of single polymorphisms during circulation in environmental, estimated by the results of genomic analysis of outbreaks in Central Africa (the phylogenetic proximity of pathogens that caused outbreaks in the DRC in 1995, 2014 and 2018 indicates the presence of a common ancestor) is 4–6×10^−4^ substitutions per site per year, which over 5-6 year period after the EVD epidemic would manifest itself in the accumulation of about 50 SNPs (**Figure 1b**). Given that the causative agent of the 2021 outbreak accumulated 10 SNPs instead of 50, the hypothesis of a environmental reservoir can be regarded extremely unlikely.

## Discussion

The main working hypothesis is currently considered the long-term persistence of the virus in certain areas and organs of the body that are not susceptible to antibodies, including the spinal cord, brain, eyeballs, placenta, mammary glands, and testicles after infection and clinical recovery [5, 6, 7]. Indeed, after the EVD epidemic of 2013–2016, it was shown that a viable virus can remain and be secreted for long periods of time, in particular, with semen in survivors up to one and a half years on [8, 9, 10]. Sexual transmission of the virus from late relapses was confirmed as the main cause of epidemiological distress in Liberia in 2015-2016 [11, 12]. Similar cases were also noted in the Republic of Guinea [13, 14]. Another typical example of the possible epidemiological significance of survivors is the case of the nurse, Pauline Cafferkey, who was infected with the Ebola virus while working at a treatment center in Sierra Leone in 2014, later evacuated and treated in the UK and was discharged with recovery [15]. It was shown that the virus remained in her cerebrospinal fluid and led to a relapse after 10 months [16]. It is of fundamental importance that the genome of the original agent and the genome isolated after 10 months from the same patient differ by only 2 single substitutions, while in the case of relay transmission from person to person, about 20 polymorphisms would be accumulated [17]. Similar frequency of SNPs occurrence in the genome of a persistent virus in the body were demonstrated in the case of EVD recurrence in DR Congo, where the period between the disease and relapse was 5 months [18]. In other word, the frequency of SNPs occurense in genom under the conditions of a survivor organism is reduced by an order of magnitude compared to that with epidemic transmission and amounts to 1–2×10^−4^ substitutions per site per year.

## Conclusion

Thus, when evaluating the hypotheses of the origin of EVD outbreak in Guinea in 2021, the most probable one should be considered infection from one of the survivor who recovered from EVD in 2014-2015.

The long period of persistence of the virus in the body of survivors and the possibility of a relapse of the disease confirms the relevance and practical significance of monitoring programs for survivors in the countries most affected by EVD outbreaks. Effective monitoring of survivors at the national level will reduce the risks of large-scale EVD outbreaks, which will significantly increase the epidemical safety of foreign countries in terms of probable importation and spread of the disease along the most loaded passenger traffic flows.

## Supporting information

will be used for the link to the fille on the preprint site

It's file contains comments to figures

## Ethical statement

Ethical approval to conduct the study was not needed. Biological specimens were collected as part of the Republice of Guinea Ministry of Health public health emergency response; therefore, consent for sample collection was waived. Personal information was anonymised in this publication.

