## Supplementary material for "Ebola virus disease outbreak in the Republic of Guinea 2021: hypotheses of origin": will be used for the link to the fille on the preprint site

Table

Nucleotide polymorphisms in genome ZE from sample E2, relatively reference Makona-Kissidougou-C15 (KJ660346, 17.03.2014, Guinea).

| POS | TYPE | REF | ALT | EVIDENCE | FTYPE | NT_POS | AA_POS | EFFECT | GENE | PRODUCT |
|---|---|---|---|---|---|---|---|---|---|---|
| 800 | snp | C | T | T:12 C:0 | CDS | 331/2220 | 111/739 | missense_variant c.331C>T p.Arg111Cys | NP | nucleoprotein |
| 1849 | snp | T | C | C:105 T:0 | CDS | 1380/2220 | 460/739 | synonymous_variant c.1380T>C p.Asp460Asp | NP | nucleoprotein |
| *2209 | snp | T | C | C:179 T:1 | CDS | 1740/2220 | 580/739 | synonymous_variant c.1740T>C p.Ser580Ser | NP | nucleoprotein |
| *2379 | snp | A | G | G:272 A:5 | CDS | 1910/2220 | 637/739 | missense_variant c.1910A>G p.Gln637Arg | NP | nucleoprotein |
| *3008 | snp | T | C | C:4 T:0 | mRNA | | | non_coding_transcript_variant | NP | nucleoprotein |
| *3050 | snp | C | T | T:6 C:0 | mRNA | | | non_coding_transcript_variant | VP35 | VP35 |
| 6283 | snp | C | T | T:16 C:0 | CDS | 245/892 | 82/296 | missense_variant c.245C>T p.Ala82Val | GP | ssGP |
| *6551 | snp | C | T | T:23 C:0 | CDS | 513/892 | 171/296 | synonymous_variant c.513C>T p.Tyr171Tyr | GP | ssGP |
| 8928 | snp | A | C | C:291 A:6 | CDS | 420/867 | 140/288 | synonymous_variant c.420A>C p.Pro140Pro | VP30 | VP30 |
| *12030 | snp | T | G | G:104 T:0 | CDS | 450/6639 | 150/2212 | missense_variant c.450T>G p.Phe150Leu | L | polymerase |
| *13011 | snp | C | T | T:28 C:0 | CDS | 1431/6639 | 477/2212 | synonymous_variant c.1431C>T p.Asp477Asp | L | polymerase |
| 15963 | snp | G | A | A:20 G:0 | CDS | 4383/6639 | 1461/2212 | synonymous_variant c.4383G>A p.Lys1461Lys | L | polymerase |
| *16298 | snp | T | C | C:11 T:10 | CDS | 4718/6639 | 1573/2213 | missense_variant c.4718T>C p.Val1573Ala | L | polymerase |
| 16514 | snp | G | A | A:27 G:0 | CDS | 4934/6639 | 1645/2212 | missense_variant c.4934G>A p.Ser1645Asn | L | polymerase |
| *16834 | snp | G | A | A:2 G:0 | CDS | 5254/6639 | 1752/2213 | missense_variant c.5254G>A p.Glu1752Lys | L | polymerase |

| | | | | | | | | | | |
|---|---|---|---|---|---|---|---|---|---|---|
| 17142 | snp | T | C | C:20 T:0 | CDS | 5562/6639 | 1854/2212 | synonymous_variant c.5562T>C p.Phe1854Phe | L | polymerase |
| *17882 | snp | A | T | T:184 A:2 | CDS | 6302/6639 | 2101/2212 | missense_variant c.6302A>T p.Asn2101Ile | L | polymerase |

*- Polimorphisms unique to the genome of the ZE virus from the E2 sample
