## Supplementary material for "Ebola virus disease outbreak in the Republic of Guinea 2021: hypotheses of origin": It's file contains comments to figures

Figure 1.a. Phylogenetic relations between Zaire ebolavirus genomes presented in the NCBI GenBank database. WA - West Africa (Sierra Leone, Republic of Guinea and Liberia, 2014-2015); DRC - Democratic Republic of the Congo (1976, 1977, 1995, 2007, 2017-2020); GAB - Gabon (1994, 1995, 2011, 2002). b .Strains from the DR Congo isolated in different time periods (from 1995 to 2018) form a separate phylogenetic cluster. The established rate of mutations is 4-6 x10-4 substitutions per site per year.

Figure 2. Phylogenetic relations between the Zaire ebolavirus genomes presented in the NCBI GenBank database from West Africa (Sierra Leone, Republic of Guinea and Liberia, 2014-2015) and the ZE genome from sample E2 (Republic of Guinea, 2021).

Figure 3. Phylogenetic relations between the Zaire ebolavirus genomes presented in the NCBI GenBank database from West African countries (Sierra Leone, Republic of Guinea and Liberia, 2014-2015), closest to the ZE genome of sample E2 (Republic of Guinea, 2021). The genome of the ZE from the E2 sample differs by 10 SNPs from the genomes of the closest group of ZE strains (MH425138, KT725387, KT725385, KT725263, KR817134, KR817132, KR817123, KR817121, KR534588) isolated from in summer and autumn 2014
